# Eco-evolutionary processes underlying early warning signals of population declines

**DOI:** 10.1101/422915

**Authors:** Gaurav Baruah, Christopher F. Clements, Arpat Ozgul

**Affiliations:** Department of Evolutionary Biology and Environmental studies, University of Zurich; School of Biological Sciences, University of Bristol

**Keywords:** stability, early warning signals, population declines, eco-evolutionary factors, trait dynamics, quantitative genetics

## Abstract

1. Environmental change can impact the stability of populations and can cause rapid declines in abundance. Abundance-based warning signals have been proposed to predict such declines, but these have been shown to have limited success, leading to the development of warning signals based on the distribution of fitness-related traits such as body size.
2. The dynamics of such traits in response to external environmental perturbations are controlled by a range of underlying factors such as reproductive rate, genetic variation, and plasticity. However, it remains unknown how such ecological and evolutionary factors affect the stability landscape of populations and the detectability of abundance and trait-based warning signals of population decline.
3. Here, we apply a trait-based demographic approach and investigate both trait and population dynamics in response to gradual changes in the environment. We explore a range of ecological and evolutionary constraints under which the stability of a population may be affected.
4. We show both analytically and with model-based simulations that strength of abundance-based early warning signals is significantly affected by ecological and evolutionary factors.
5. Finally, we show that a unified approach, combining trait- and abundance-based information, significantly improves our ability to predict population declines. Our study suggests that the inclusion of trait dynamic information alongside generic warning signals should provide more accurate forecasts of the future state of biological systems.

## 1. Introduction

Predicting catastrophic and non-catastrophic collapses of populations and ecosystems in response to novel environmental pressures are critical if we are to mitigate the effects of global change and minimize the loss of biodiversity (Cardillo et al., 2005; Thomas et al., 2004). These collapses are however difficult to predict (Boettiger & Hastings, 2012b; Clements, Drake, Jason, & Ozgul, 2015; Clements & Ozgul, 2018). One suite of methods, which may achieve this, are early warning signals derived from dynamic systems theory (Scheffer et al., 2009b; van Nes & Scheffer, 2007). Such methods assume that a dynamical system, for example a biological population, that is under increasing continuous external pressure, will show a property called critical slowing down (CSD) as it approaches a tipping point-the point at which the system’s state can substantially change in response to a slight perturbation. Around the vicinity of this tipping point, the system takes longer to return to its original state after every perturbation (Wissel, 1984; Strogatz, 1994; Scheffer *et al.*, 2009).

The occurrence of CSD is a direct implication of the dominant eigenvalue of the system approaching zero (for continuous dynamical system) (van Nes & Scheffer, 2007) or 1 (for a discrete dynamical system) (Krkoek & Drake, 2014). As a direct consequence of CSD, statistical moments embedded within abundance time-series data, particularly variance and autocorrelation will show marked increase over time. These two indicators are the prominent abundance-based statistical metrics, and, along with various other temporal (Boettiger & Hastings, 2012a; Chevalier & Grenouillet, 2018; Vasilis Dakos et al., 2012; Drake & Griffen, 2010; Jarvis, McCann, Tunney, Gellner, & Fryxell, 2016) and spatial (Butitta, Carpenter, Loken, Pace, & Stanley, 2017; Dai, Korolev, & Gore, 2013) metrics, are commonly known as early warning signals (EWS).

Changes in autocorrelation and variance are measures of the stability of a system, derived from the theory of alternative stable states (Clements & Ozgul, 2018; Wissel, 1984). Holling’s (1973) fundamental work on stability and resilience laid the foundation for future work on deriving stability criteria for systems with alternative equilibrium states. Stability of a dynamical system can be characterized by the classic ball-in-cup diagram (Clements & Ozgul, 2018; Holling, 1973; Nolting & Abbott, 2015), where the ‘ball’ represents the state of the system and the ‘cup’ represents the potential energy landscape (Fig.1). The stability of the system is thus quantified by the depth of the energy landscape, and resilience by the size of the cup (Clements & Ozgul, 2018; Villa Martín, Bonachela, Levin, & Muñoz, 2015). In response to continuous exogenous pressure, the system loses resilience (for example, decline in population numbers) and the energy landscape shrinks (Fig. 1). Concurrent with the shrinkage of the landscape, there is also a decrease in the depth of the ‘ball-in-cup’ landscape signaling a loss in stability (Fig. 1). It is this loss in stability, which statistical EWSs such as variance and autocorrelation measure as an alternative for loss of global resilience of the system (Clements and Ozgul 2018). The shape of the ball-in-cup landscape thus determines the dynamics of the state variable in response to an exogenous environmental change (Beisner, Haydon, & Cuddington, 2003).

**Figure 1.**
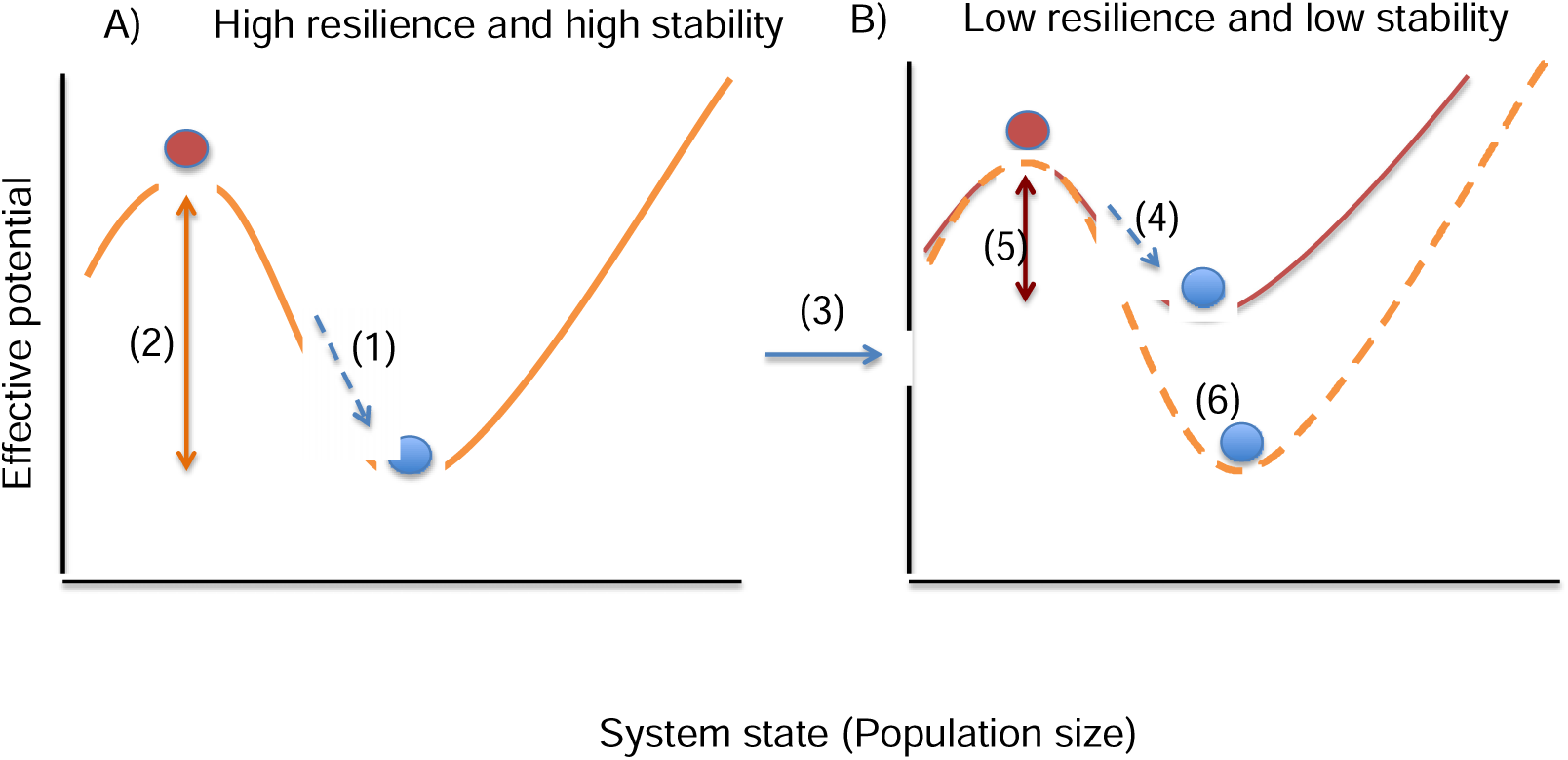
Illustration of the ‘ball-in-cup’ potential energy curve in relation to stability and resilience of a one-dimensional dynamical system. The stable state of the dynamical system (A) which is an isolated population, can be indicated by the blue ball, and the unstable state (or the tipping point) can be indicated by the red ball in both A and B. (1)-shows that the slope of the potential curve (i.e., stability) is steep and the basin where the ball lies is also wide signifying that both resilience as well as stability is high. This means that the amount of effective potential energy needed (2) to push the ball from its stable state (blue) to the unstable state (red) (or in other words the reduction in stability) will be large. However, for another isolated population, there might be intrinsic differences in ecological or evolutionary factors controlling the dynamics of that isolated population and this might cause (3) a shift in the ‘ball-in-cup’ potential energy curve from (6) to (4), for that population. In (4) the slope is now less steep, signifying a reduction in intrinsic stability. Along with this, there is also a reduction in intrinsic resilience. Now, the amount of energy (5) to move the blue ball at (4) to the unstable state for (B) is less. This could reflect in the performances of EWS of population declines.

However, the shape of the ‘ball-in-cup’ landscape can be changed not only by exogenous pressures, but can also be altered by intrinsic factors associated with the system’s dynamics (Beisner et al., 2003). Changes in these intrinsic factors might alter the location of an equilibrium state, or destabilize the current state, causing the system to arrive at a point closer to the other alternative state (Fig. 1). For a biological population, the state of the population will be characterized by its current abundance. Responses of this state variable (abundance) to an exogenous environmental pressure will be a combination of ecological and evolutionary responses controlled by factors such as genetic variation or plasticity (Ozgul et al., 2010; Price, Qvarnström, & Irwin, 2003). For example, plastic responses to fluctuating changes in the exogenous environment are usually fast which consequently stabilizes population fluctuations (Reed, Waples, Schindler, Hard, & Kinnison, 2010). However, if the environment keeps on changing, the population might not only deplete its plastic capacity but also its standing genetic variation, which in turn leads to its eventual decline. Further, such a decline would also be conditional on the population’s reproductive rate, with higher reproductive rate leading to a slower decline (Juan-Jordá, Mosqueira, Freire, & Dulvy, 2015). In addition, due to continuous changes in the exogenous environment, selection pressures can subsequently increase leading to contemporary evolution (Yoshida, Jones, Ellner, Fussmann, & Hairston, 2003). Adaptive evolution would then depend on the amount of genetic variation present in the population with higher variation leading to faster evolution, which can either stabilize or destabilize population dynamics (Cortez, 2016; Sanchez & Gore, 2013). Since the dynamics of a population in response to an exogenous pressure is determined by the shape of its potential energy landscape (Beisner et al., 2003), it is thus expected that these factors namely plasticity, genetic variation, and reproductive rate have the capacity to influence the shape of the ball-in-cup landscape. In turn, given that EWS are stability measures derived from alternative stable states theory (Beisner et al., 2003; Clements & Ozgul, 2018; Scheffer et al., 2009a), it is very likely that strength in EWS and hence predictability of population decline might also be affected by these factors. Such theoretical expectations raise important practical questions: can predictability of population decline with the help of EWS be affected by such ecological and evolutionary factors?

Abundance-based measures of the stability of an ecological system might not be the only indicators of a population being forced by an external environment (Clements & Ozgul, 2016b). Phenotypic traits, such as body size, is one of the important traits that determines the fate of individuals as well as dynamics of population, and thus can provide crucial information regarding the current state of the population (Clements and Ozgul 2016b; Ozgul et al. 2014; Spanbauer et al. 2016). Recently, Clements & Ozgul (2016) showed, using data from experimental microcosms, that including body size information with EWS could significantly improve the predictive accuracy of population declines. Such trait-based signals, have been further shown to be present in the historic collapses of whale populations (Clements et al., 2017). However, extensive research on the efficacy of such signals in comparison to abundance-based EWS is still in its infancy (Clements & Ozgul, 2018).

Here in this paper, we used a quantitative genetics framework to simulate the dynamics of a population in response to gradual environmental change. We track both the population dynamics and trait distribution until the simulated population declines significantly. We show that this particular model exhibits critical slowing down behavior and hence is expected to show EWS. We then assess whether genetic variation, plasticity in the trait, and net reproductive rate alter the ball-in-cup landscape (stability landscape) and consequently the strength in EWS. In addition, we assess the efficacy of trait-based EWS in comparison with abundance-based EWS and reiterate the ability of trait-based signals in accurately informing population declines.

## 2. Materials and Methods

### 2.1 Mathematical model

We model the joint dynamics of quantitative trait evolution and population size under density dependent regulation via the effects of selection (Chevin, Lande, & Mace, 2010; Chevin & Lande, 2010; Gomulkiewicz & Holt, 1995). We consider a population that has discrete generations where individuals have fitness that is determined by a single quantitative trait z under stabilizing selection with linear reaction norms (Gavrilets & Scheiner, 1993). Under these assumptions the dynamics of the population and the mean value of the trait can be written as (Lande, 2009):

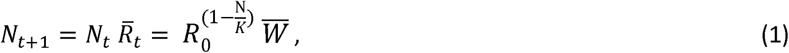

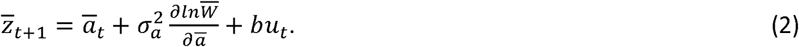

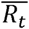 is the growth rate of the population at generation t, R_0_ is the net reproductive rate which is under density dependent selection given by the exponent 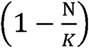(Chevin & Lande, 2010). The average fitness of the population is given by the equation 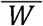 which is due to the quantitative trait *z*.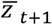 is the average value of the trait at generation *t+1*, 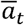 is the mean breeding value or genotypic value of the population at generation *t*, 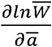 is the gradient of selection on the mean trait 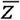, 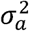 is the additive genetic variance,*bu*_*t*_ quantifies the average plastic response of the trait at time *t*, and *u* is the environmental cue (table 1).

Expanding 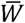 in equation (1) as follows:

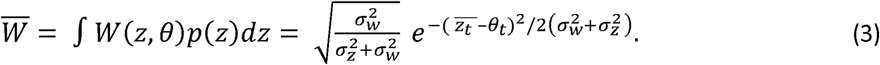

**Table 1:** List of parameter values and variables used in the model simulations. The variables are not given any value but factors for which the model simulations are tested are given values.

| Parameters/variables | Description | Value |
| --- | --- | --- |
| $N_t$ | Abundance of the population at time $t$ | .... |
| $\bar{z}_t ; \bar{a}_t$ | Mean trait value at time $t$ ;<br>mean breeding value at time $t$ | .... |
| $\sigma_z^2 ; \sigma_a^2$ | Variance of the mean trait ;<br>additive genetic variance or<br>variance of the mean<br>breeding value | [0.05, 0.1, 0.2, 0.3, 0.4,0.5] ;<br>[0.05,0.1, 0.2, 0.3, 0.4,0.5] |
| $b$ | Strength in phenotypic<br>plasticity | [0.05, 0.1, 0.3,0.4, 0.8] |
| $R_0$ | Net reproductive rate | [1.1, 1.2,1.3,1.4, 1.5] |
| $u_t$ | Environmental cue at time $t$ | $U_t = \begin{cases} 0 & , t < 500 \\ 1 - c \frac{t}{500} & , t \geq 500 \end{cases}$ |
| $\theta_t$ | Optimum phenotype at time $t$ | $\theta_t = BE_t = \begin{cases} 0 & , t < 500 \\ B - B \frac{t}{500} & , t \geq 500 \end{cases}$<br>($B = 20$) |
| $w_z^2$ | Width of the Gaussian<br>stabilizing fitness function. A<br>measure of the strength in<br>selection | 20 |
| $z_i, a_i$ | Trait value for an individual $i$<br>; breeding value for an<br>individual $i$ | .... |
| $\epsilon$ | Residual component or<br>unexplained component of a<br>trait $z$ | 0 |
| $\sigma_\epsilon^2$ | Variance in that residual<br>component $\epsilon$ | 0 |
| $E_t$ | External environmental value<br>at time $t$ | .... |
| $B$ | Coefficient that captures the<br>strength in the external<br>environmental change | 20 |
| $c$ | Coefficient that controls the<br>rate of change in the<br>environmental cue | 5 |
| $\xi$ | Environmental predictability; | 1 |
|  | quantifies how much the external environment is correlated with the environmental cue |  |
| $\sigma_\theta, \sigma_u$ | Variance in optimum phenotype, variance in environmental cue | $1.25^2, 1.25^2$ |

The integral in (3) over all trait values in the population gives the average fitness of the population *p*(*z*) is the distribution of the trait in the population, 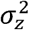 is the variance in the phenotypic distribution *p(z)* and *W*(*z, θ*) is the gaussian stabilizing fitness function given as:

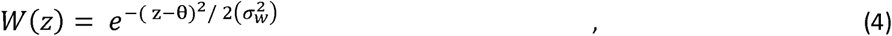

with width 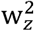 and optimum phenotype of *θ*. Individuals with trait values far from the optimum will have lower fitness compared to individuals whose trait value matches the optimum phenotype *θ*. Hence an individual’s fitness will be determined by how far its trait value *z* is from the optimum phenotype *θ*, (Chevin and Lande, 2010). Finally, the response of the primary phenotype *z* of an individual in this finite population to the external environment is modeled using linear reaction norms (Gavrilets and Scheiner, 1993; Lande and Shannon, 1996):

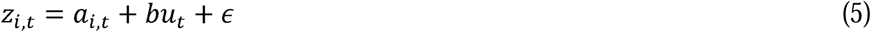

where *a* is the breeding value of the individual *i*. The breeding value is normally distributed with mean 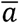 and additive genetic variance 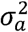, and *ϵ* is the residual component with mean 0 and variance 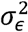. The slope *b* quantifies how plastic the trait *z* is in response to the environment. In our model *b* is constant meaning that plasticity in the trait cannot evolve (Chevin & Lande, 2010). We model an environment and the environmental cue (see below) that determines the optimal phenotypic value *θ*_*t*_ for the primary phenotype *z*. The optimal phenotype *θ*_*t*_, is assumed to be linearly dependent on the external environment *E* that selects for a particular phenotypic value such that,

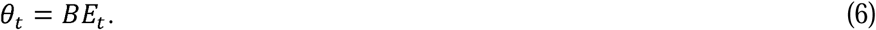

Expanding equation 6, the dynamics of the optimal phenotype is:

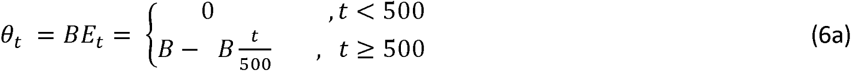

And the environmental cue *u* in equation (5) is modeled as:

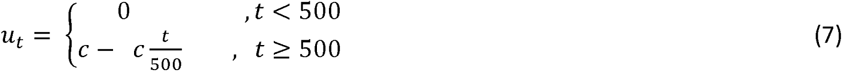

Environmental change is modeled at t = 500, where a shift in the external environmental value *E_t_* also triggers a shift in the optimum phenotypic value. The rate of the shift in the optimum phenotypic value is controlled by the parameter B (equation 6a, see section 2.4 below). As in Reed et al. (2010), the dynamics of the optimal phenotype *θ*, in equation (6a) and the environmental cue *u* in equation (7) are modelled to be correlated over time. This correlation is modeled by taking the help of a bivariate normal distribution: at each generation *t,* a value for the optimal phenotype *θ* and environmental cue *u* are drawn randomly from a bivariate normal distribution with means given by equation (6a) and (7). The covariance of the bivariate normal distribution is given by *σ*_*θ,u*_ = *ξ σ*_*θ*_*σ*_*u*_ where *ξ* is the correlation between *θ* and the cue *u*. If *ξ* = 1, then the values of the optimum phenotype and the environmental cue drawn from the bivariate normal distribution will be perfectly correlated over time *t* and when *ξ* = 0, they will not be correlated. Thus, determines predictability of the external environment such that = 1 means that an individual is able to perceive changes in the external environmental perfectly with the help of an environmental cue *u* that is correlated with the changes in the external environment. In such a case, the plastic response will track the changes in the environment and the strength of the plastic response of the trait will be determined by *b*. We assumed that variances in the cue *σ*_*u*_ and the optimum *σ*_*u*_ are equal (see table 1). This mimics the situation, for instance, when variation in food availability is correlated with inter annual variation in snow melt (Vuren & Armitage, 1991) - where in this case, inter annual variation of snow melt is the environmental cue *u* that the organism perceives, and variation in food availability is the external environment *E*. Depending on the changes in the external environment (for example, changes in food availability), the optimum phenotype *θ*, will change which is given by equation 6.

The above model is based on the infinitesimal quantitative genetic model (Chevin & Lande, 2010; Falconer & Mackay, 1996).

### 2.2 Stability landscape and effective potential

One approach to quantifying stability which is also fundamental to dynamical systems with alternative stable states is the ‘ball-in-cup’ analogy (Haydon, 2003; Villa Martín et al., 2015). The different states of the dynamical system can be represented by a landscape and the state of the population at a particular moment by a ball in that landscape. The landscape of the dynamical system can be represented by an effective potential *V*(*n*) (Beisner et al., 2003; Villa Martín et al., 2015). If a dynamical system (example a population) can be represented by a continuous differential equation in the form of: 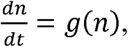, then the effective potential of that dynamical system can be written as : *V*(*n*) = − ∫ *g*(*n*)*dn* (Beisner et al., 2003; Nolting & Abbott, 2015). For our dynamical system, 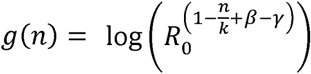 (see appendix 4). Here *n* is the state of the system, which in our case is the log of the population size *N*. The effective potential of our dynamical system is then (appendix 4):

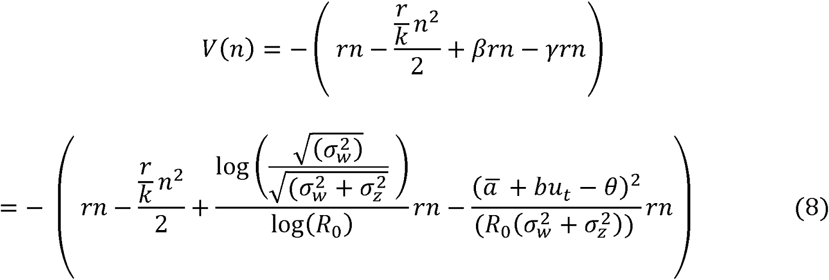

It is evident from the above equation (8) that the effective potential function is dependent on three parameters: *r*, *β*, *γ*, where 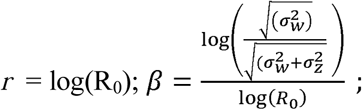; 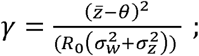; *n* = log(N); kμ log(K). *γ* represents the standing load of the population and represents the lag-load.

### 2.3 Equilibria and eigenvalues of the model

The extinction equilibrium for the model is at *N* = 0* and the positive equilibrium for the model is at N* = *K* + *γK* − *γK*, where *γ* is the lag-load and represents the standing load of the population (see appendix 1). Stability of the two equilibria are based on their corresponding eigenvalues, 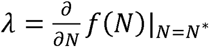. Equilibria will be stable if magnitude of *λ* < 1 and unstable if magnitude of *λ* > 1. For details of linear stability analysis and occurrence of CSD see appendix 1 and appendix 2.

### 2.4 Simulations of population declines

We performed stochastic simulations of the model described in section 2.1. Dynamics of the trait, population and the external environment were iteratively updated using equations (1-7). Without loss of generality, the mean phenotypic value 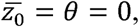, at the start of the simulation, i.e., at t=0 (equation 6a). This meant that initially before a shift in the external environment, average phenotypic value of the population is at its optimum. External environmental change in our model was enforced at t=500 after the populations had reached the carrying capacity. Stochasticity in the model was introduced at each generation *t* by randomly drawing both optimum phenotypic value and the environmental cue from a bivariate normal distribution with means given by equation 6a and 7 respectively, and covariance given by *σ*_*θ,u*_ = *ξ σ*_*θ*_*σ*_*u*_, where *σ*_*θ*_ and *σ*_*u*_ quantifies the variance in the fluctuation of optimum phenotype and the environmental cue over time *t*. Environmental predictability in our model was set at *ξ* = 1, which meant that the external environmental change and the environmental cue drawn at each generation *t* were highly correlated (see Reed et al. 2010). We forced the populations to decline by changing the rate of change in the environment E_t_, at t = 500. We chose a rate of change in the environment in a way that populations declined regardless of any combination of parameter values. This was done by setting B =20 in (6a) in all of our simulations, after *t*≥500. In all of our simulations, for each parameter value of a factor (genetic variation, plasticity, net reproductive rate) we did 100 independent simulations of population declines.

#### 2.4.1 Abundance-based EWS

For abundance-based EWS analyses, the length of the time-series used was of the same length for each factor (genetic variation, plasticity, net reproductive rate) in question. For each parameter value of each factor in question, we used 30 time-series data after the time point of environmental change i.e., after t= 500 to t=530. We discarded the rest of time-series data. We chose a rate of environmental change in a way (B=20) that population decline was not immediate. Significant population declines were observed across replicates for different factors after t = 530 time point (see timeseries data, Fig. S3-S5). Next, we analyzed 30 abundance data points after t=500, when the environmental change sets in. This ensured normalization of the length of each time series and minimized the effect of varying length of timeseries. We then applied generic EWS of population collapse by using the *earlywarnings* package in R. We focused our analysis on two specific indicators namely autocorrelation at the first-lag (AR1) and standard deviation (SD) as most of the other indicators – such as first-order autoregressive coefficient or coefficient of variation – could be theoretically derived from these parameters (Dakos et al. 2012). These two EWSs are theorized to increase over time before a significant population decline (Dakos, Carpenter, van Nes, & Scheffer, 2014; Scheffer et al., 2009a). Thus, the important question is whether these two EWSs increases before significant population declines that was observed after time point t = 530 (see Fig. S3-5). If the two EWSs increased over time before significant population declines, were their performances affected by the different ecological and evolutionary factors?

The two indicators AR1 and SD were calculated using a predefined sliding window which was 50% of the abundance time-series being analyzed. We z-standardized the leading indicators so that it would be easier to incorporate information from phenotypic trait later in our analyses. We used Gaussian detrending to remove any trend in the abundance time-series and used Kendall’s Tau correlation coefficient of the indicators prior to time point t=530 as an indicator of an approaching decline. Kendall’s tau is a non-parametric measure of rank correlation used to identify an increasing trend. Kendall’s tau takes values in the range of [-1,1]. Strong positive Kendall’s Tau correlations of the statistical indicators (SD and AR1) with time would indicate an approaching population decline (Dakos et al. 2012). If Kendall’s tau correlation coefficient for an EWS was less than 0.5, we considered that EWS as a false negative. Hence, we quantified reliability of EWS as the rate of false negatives: the number of times in replicate simulations the Kendall’s tau correlation coefficient was less than 0.50. Higher proportion of false negatives would indicate less reliability of EWS and vice-versa.

#### 2.4.2 Trait-based EWS

To evaluate the efficacy of trait-based EWS (Clements & Ozgul, 2016b), information from mean phenotype (was incorporated with the leading indicators in an additive manner (Clements *et al.*, 2017; Clements and Ozgul, 2018). To include information from average phenotypic value, we first z-standardized the trait time-series. The length of the phenotypic time-series that was used was the same as the corresponding abundance time-series. Before a population decline, SD and AR1 are expected to increase over time, while in our model, because of the direction of the optimum environmental change, the trait is expected to decline (it is also possible for trait to increase over time, but that would depend on the direction of the optimum phenotypic change). Such a phenotypic trait, for example, could be body size. Body size is a quantitative trait fundamental to life-history theory and population dynamics (Cameron, O’Sullivan, Reynolds, Piertney, & Benton, 2013; Gibert, Allen, Hruska, & DeLong, 2017; Pigeon, Ezard, Festa-Bianchet, Coltman, & Pelletier, 2017), which has been shown to decline in response to decreases in resource availability (Clements & Ozgul, 2016b), increases in temperature (Forster, Hirst, & Atkinson, 2011, 2012; Gardner, Peters, Kearney, Joseph, & Heinsohn, 2011; Tseng et al., 2018), and increases in harvest rate (Frank *et al.*, 2011; Clements *et al.*, 2017). Whether including trait information improves predictability of population decline will be dependent on whether the trait responds to an external environmental change. Here, in our model, the optimum environmental change will affect the dynamics of the phenotype. However, whether the change in the dynamics of the phenotype will occur within the time series length being analyzed here and hence be informative of population decline is unknown. It has been shown that it is possible for shifts in phenotypic dynamics to precede population declines, but only under certain environmental circumstances (Baruah *et al.*, 2018 *(in press)*). As of yet, extensive analysis of the efficacy of EWS and trait-based EWSs has not been done.

Since the phenotype is expected to decrease in response to the change in the optimum environment, the z-standardized values of phenotypic timeseries were multiplied by - 1 so that they could be included with the other two standardized statistical metrics such as SD and AR1. The values of standardized EWS such as SD and AR1 were added additively with standardized mean phenotypic time series to quantitatively create trait-based signals or trait-based EWS. We evaluated three trait-based EWS metrics namely *AR1+trait, AR1+SD+trait, SD+trait* and compared how the metrics that included information from the mean phenotype improved predictability of population declines when compared with the abundance-based EWS like *AR1, SD.* We quantified Kendall’s Tau correlation coefficient as an indicator for strength of predictability of population declines and reliability of the trait-based EWS as the rate of false negatives.

#### 2.4.3 Adaptive plasticity

To assess the role of adaptive plasticity *b*, we simulated 100 replicate population declines for each parameter value of *b,* which ranged from low to high: 0.05, 0.1, 0.3, 0.4, and 0.8. While we varied plasticity, we fixed 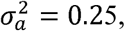, and R_0_ at 1.2. The width of the fitness function determines the strength of selection in the trait. We fixed the width of the fitness function at 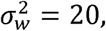,which would indicate medium strength in selection. Usually, 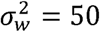 would indicate weak strength in selection and 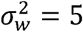 would indicate strong selection (Chevin and Lande 2010).

#### 2.4.4 Genetic variation

The levels of genetic variation 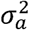 used for the simulations of population declines are 0.05, 0.1, 0.2, 0.3, 0.4, 0.5. Adaptive plasticity *b*, net reproductive rate R_0_ and 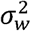 were kept at 0.2, 1.2 and 20 respectively. We also did simulations where we relaxed the assumption of genetic variation 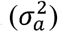 remaining constant during population collapses. For details see appendix 5.

#### 2.4.5 Net reproductive rate

To assess the role R_0_, we simulated 100 replicate population declines. The levels of R_0_ used for the simulations of population collapse were 1.1, 1.2, 1.3, 1.4, and 1.5. Adaptive plasticity *b*, genetic variation 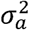 and 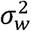, were kept at 0.2, 0.25 and 20 respectively.

## 3. Results

### 3.1 Effective potential

#### 3.1.1 Strength of plasticity

The term 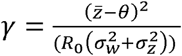 in equation (8) is the evolutionary lag due to maladaptation of the mean phenotype with the optimum (Lande, 2009). The effective potential becomes narrower with increasing strength of plasticity and consequentially increases stability, i.e., the potential function becomes more negative (Fig. 2). Since return time and stability are inversely related (Dai et al., 2015), return time decreases with increasing strength of plasticity (Appendix 2, Fig. S1). In other words, the population quickly recovers with every perturbation if strength of plasticity is high.

**Figure 2.**
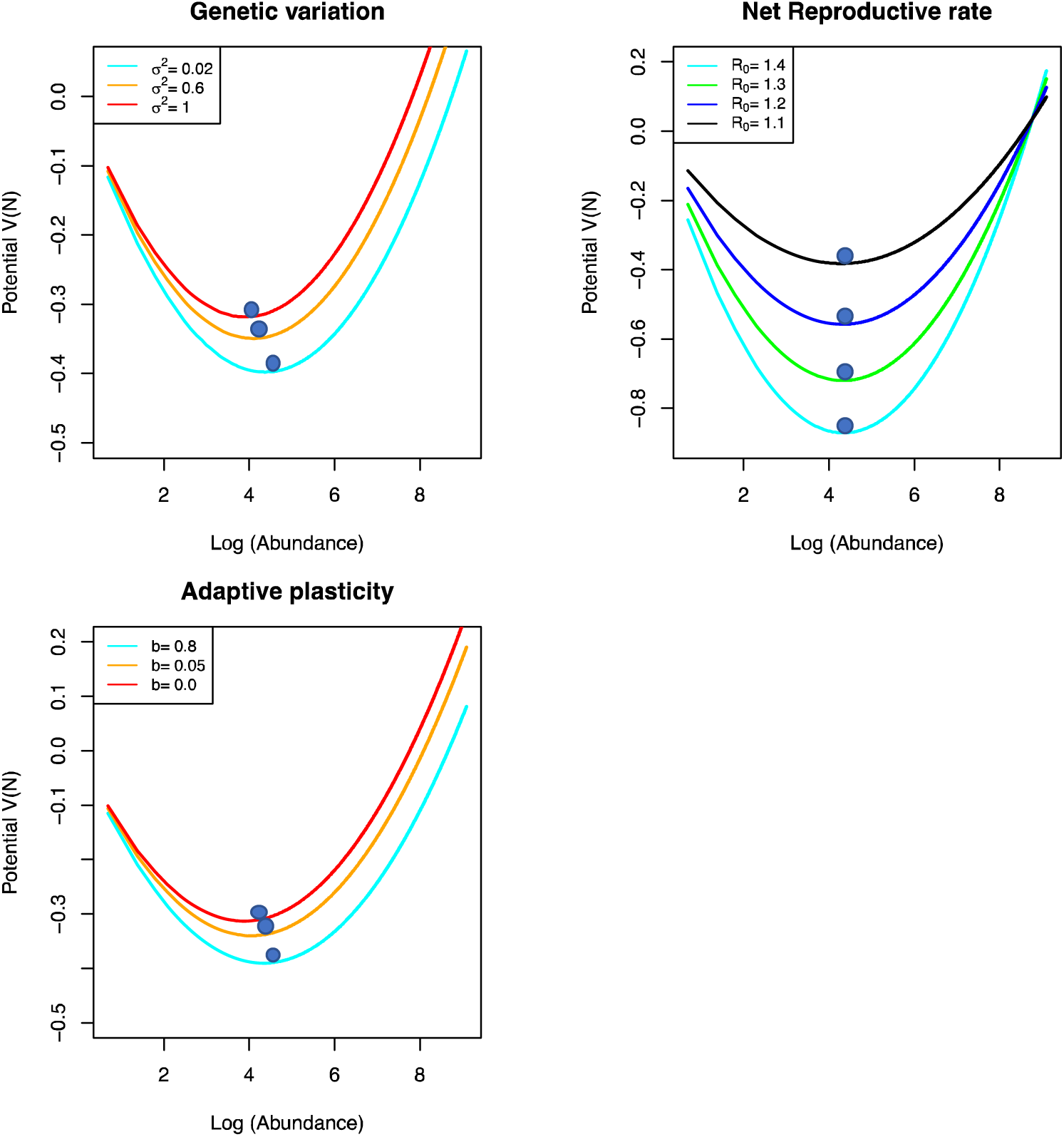
Effective potential *V(N)* plotted as a function of the state variable abundance, N. Note the effective potential is altered by factors like genetic variation, adaptive plasticity and net reproductive rate. The lower the value of the effective potential, more stable the system is.

#### 3.1.2 Genetic variation

The mean phenotype has variance of 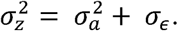. Assuming residual component *σ*_*ϵ*_ to be zero, phenotypic variance is then equal to additive genetic variance,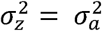. Inspection of the equation (8) reveals that high additive genetic variance causes the effective potential to be more positive when compared with low 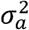 and consequently decreases stability, i.e., the effective potential becomes less negative (Fig. 2; the more negative the effective potential, more stable the population is). Reduction of stability and effective potential energy is higher for populations with low genetic variance whenever they are forced to collapse compared to populations with high genetic variance (Fig. 2). Further, return time also increases with increasing additive genetic variance (Appendix 2, Fig. S1).

#### 3.1.3 Net reproductive rate

From equation (8), we also see that higher values of R_0_ lead to higher stability and narrow potential curve for a given state of the population (Fig. 2). Loss in stability is more when R_0_ is higher for the same rate of change in the environmental change when compared with a lower R_0_. Further, return time also decreases with increasing R_0_ (appendix 2, Fig. S1).

### 3.2 Simulations of population declines: abundance-based EWS

#### 3.2.1 Strength of plasticity

The strength of EWSs of population collapse (Kendall’s Tau value) was much higher in populations which had high adaptive plasticity ***b** =* 0.8 (Fig. 3A, B) when compared with populations with low adaptive plasticity. Moreover, the rate of false negatives decreased for all the abundance and trait-based metrics as strength of plasticity increased (Fig. 5A). Rate of false negatives however remained more or less same for AR1 across different plasticity levels (Fig. 5A).

**Figure 3.**
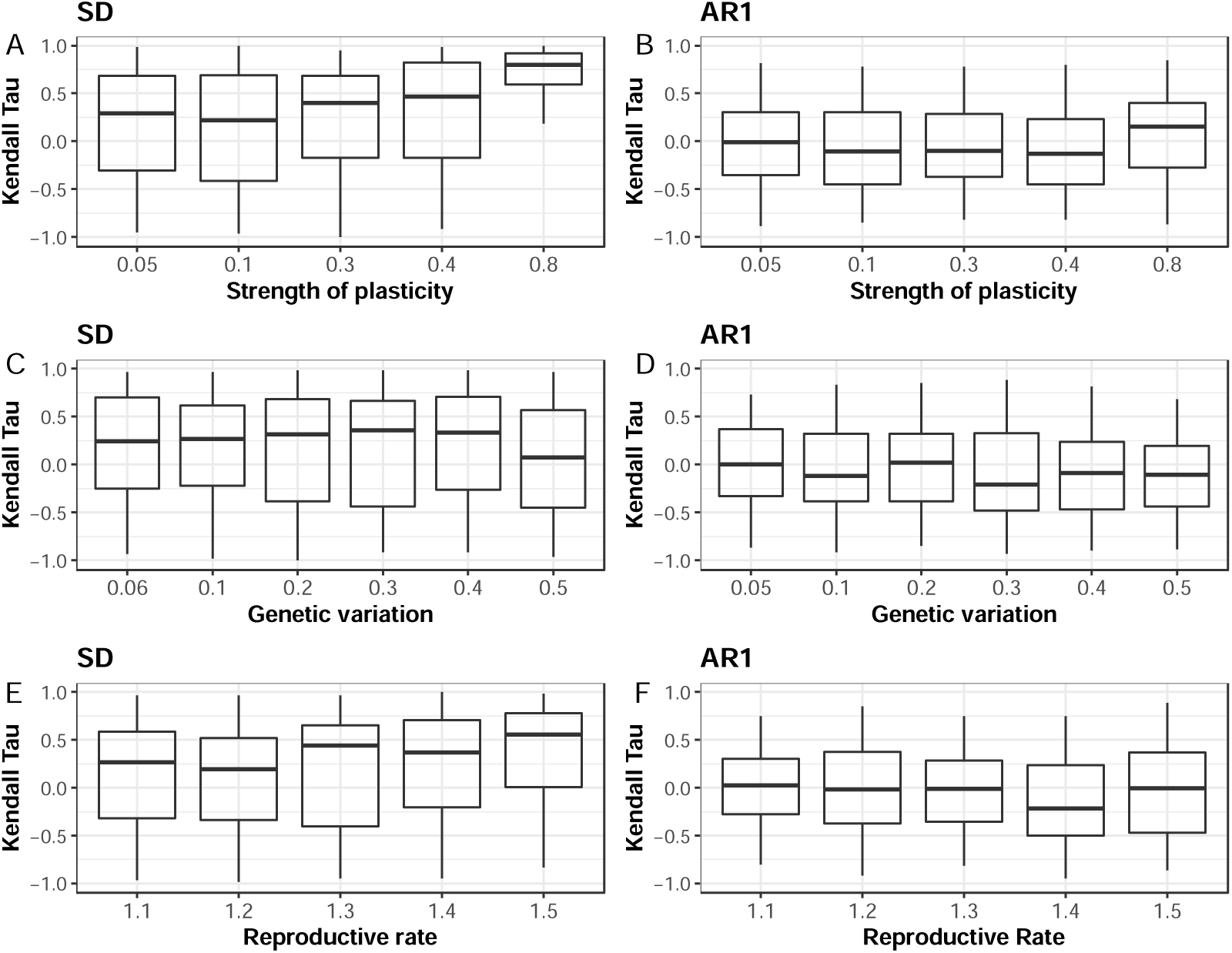
Box plots showing the distribution of Kendall’s Tau correlation coefficient (high values around 0.5 are usually considered an EWS) for 100 simulation of population declines for different levels of adaptive plasticity *b,* genetic variation 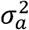, reproductive rate R_0_. The leading indicators that are shown here are autocorrelation at lag-1 (AR1) and standard deviation (SD). X-axis is the different levels of the above factors.

#### 3.2.2 Genetic variation

Higher genetic variation in the population led to slight decrease in detectability of population decline than in populations with lower genetic variation, particularly for SD (Fig. 3C, D). Moreover, the rate of false negatives for SD, AR1 remained more or less same as genetic variation decreased (Fig. 5B). For trait-based EWSs, particularly for *SD+trait*, rate of false negatives was the lowest (Fig. 5B).

#### 3.2.3 Net reproductive rate

Strength in EWS was lower in populations with lower R_0_ when compared with higher R_0_, particularly for SD (Fig. 3E). AR1 however showed a slightly different result (Fig. 3F). In case of reliability of abundance-based EWS, the rate of false negatives decreased for SD as net reproductive rate increased (Fig. 5C). However, for AR1 rate of false negatives remained unchanged as R_0_ increased (Fig. 5C).

#### 3.3 Simulations of population declines: trait-based EWS

Trait-based EWS improved the strength of EWS of population declines (Fig. 4, Fig S8-S9). In principle, *AR1* was the least efficient of all the signals compared. However, when trait information was included with *AR1*, strength in trait-based *AR1 (trait+AR1)* improved significantly (Fig.4, Fig. 5). Overall, the Kendall’s Tau value of SD and *SD+trait* was relatively higher than all the other trait-based and abundance-based EWS. This was consistent across all the factors analyzed such as genetic variation, plasticity (Fig.4, Fig. S8-S9). The rate of false negatives was highest for AR1 across all factors analyzed (Fig. 5A-C). For plasticity, the rate of false negatives decreased as strength of plasticity increased, but this was however opposite for genetic variation where the false negatives slightly increased as genetic variance increased (Fig. 5B). Furthermore, even though AR1 had the highest number of false negatives, false negatives of *AR1+trait* significantly decreased (Fig. 5).

**Figure 4.**
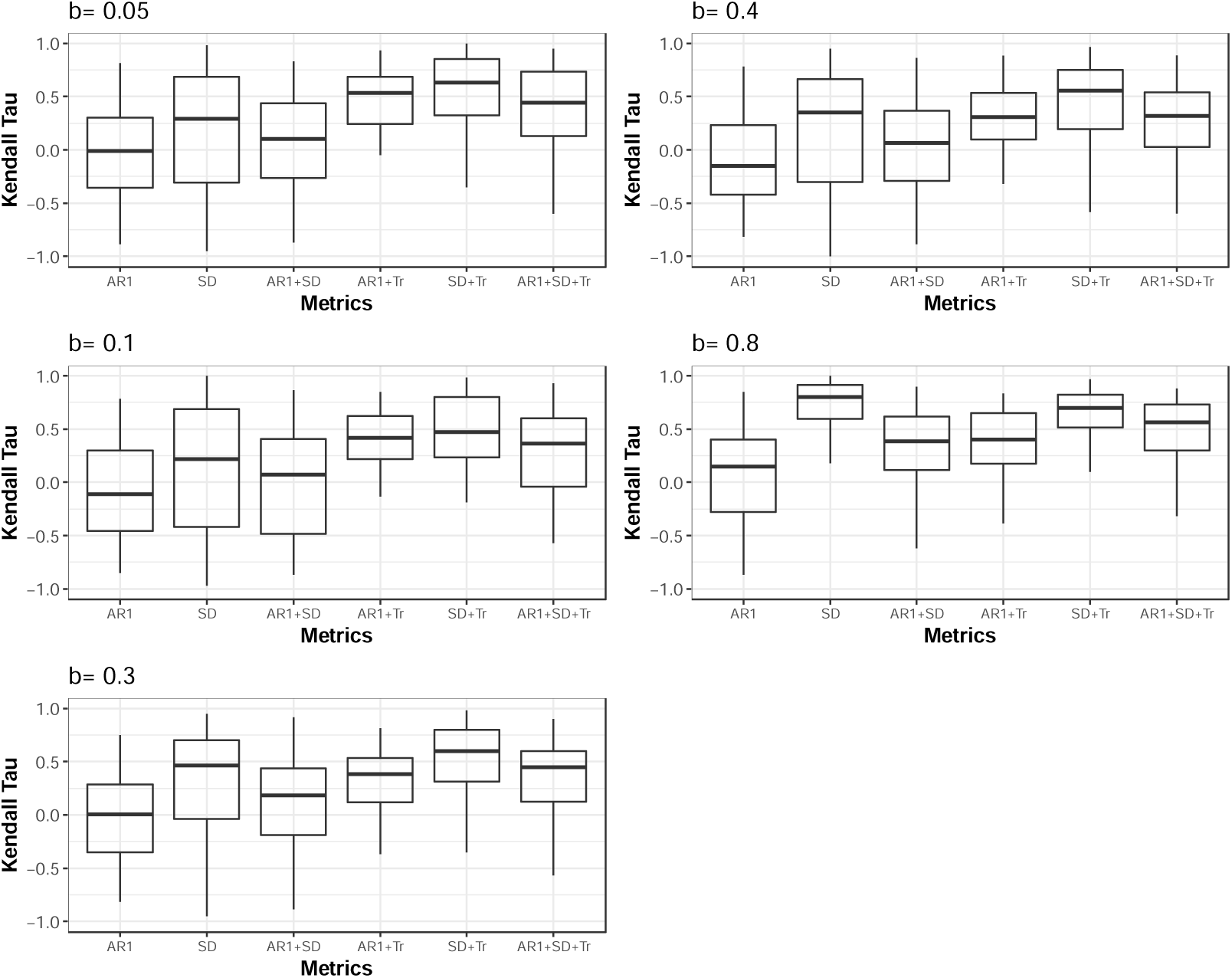
Box plots showing the distribution of Kendall’s Tau correlation coefficient for 100 simulation of population declines for different levels of adaptive plasticity *b*. On the X-axis are the different early warning metrics shown: autocorrelation at lag-1 (AR1), standard deviation (SD), *(AR1+SD),* (*AR1+Tr), (SD+Tr), (AR1+SD+Tr)*. *Tr* corresponds to the mean value of the phenotypic trait. Colored boxplots denote the trait-based EWS while white boxplots denote abundance-based EWS.

**Figure 5.**
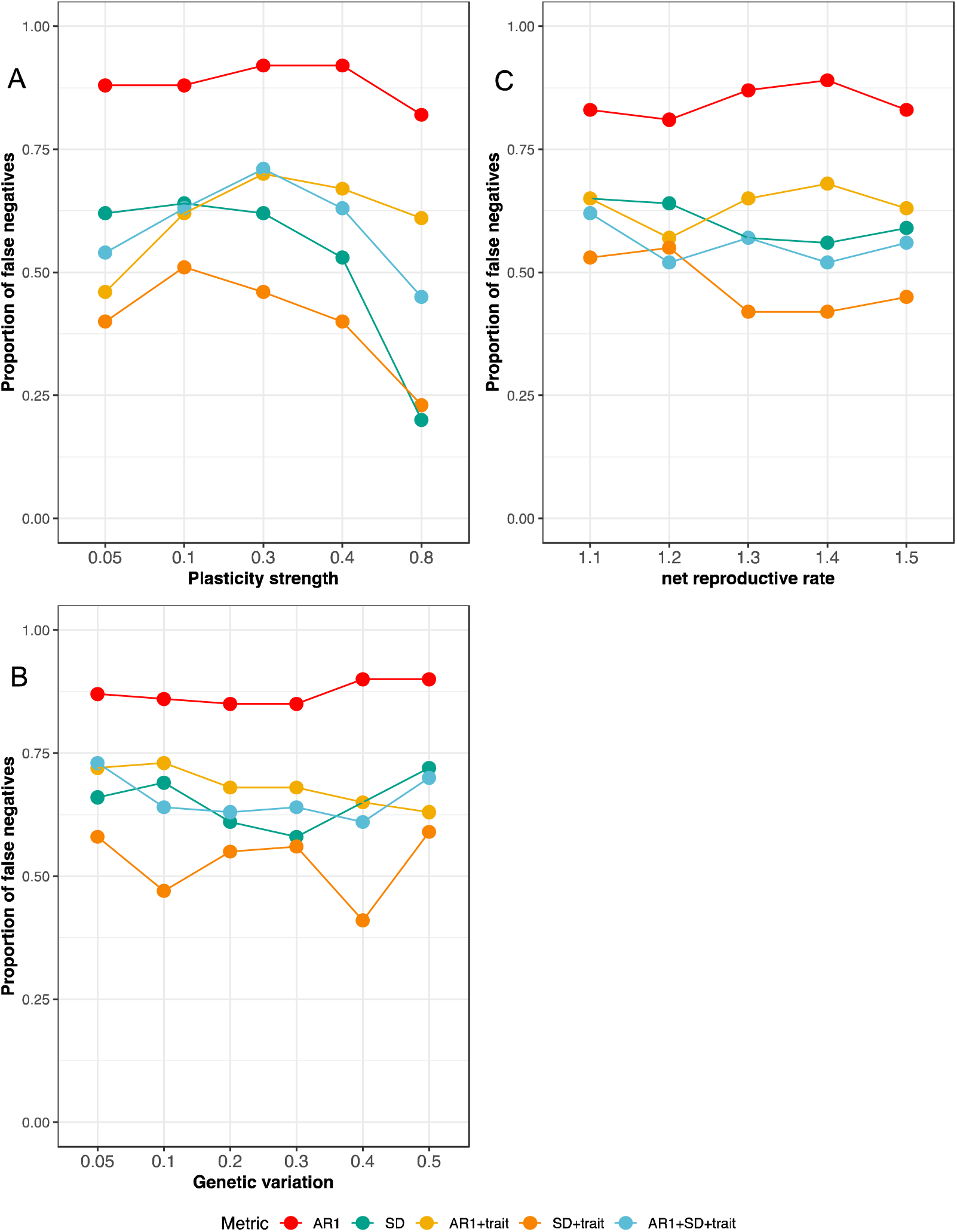
Proportion of false negatives shown here for a threshold of 0.5 (which means Kendall’s tau value below 0.5 are considered not an EWS) on the Y-axis for abundance and trait-based EWS metrics for different levels of (A) plasticity strength, (B) genetic variation, (C) net reproductive rate.

## 4. Discussion

The early detection of population declines with the help of EWS provides a unique tool to highlight conservation efforts to a population approaching an impending transition. We show in this paper that factors intrinsic to a population in question such as genetic variation, plasticity or net reproductive rate could significantly alter stability of a population as well as strength and reliability of EWS of population decline.

The detectability of EWSs is dependent not only on the relation between stability and resilience but is also determined by a variety of factors including sampling frequency, and the strength and type of external forcing (Dai, Korolev, and Gore 2015; Clements et al. 2015a; Clements and Ozgul 2016). Here we add to this body of knowledge by demonstrating that the stability of a system can also be altered by intrinsic factors such as genetic variation, plasticity and net reproductive rate. These changes to the effective potential in turn led to stronger EWS being observed, whenever changes in the external optimum caused a population to lose a significant amount of effective potential energy (Fig. 1). In other words, stronger EWSs would be observed whenever a population is forced to move from a deeper point in the potential energy curve (more stable) to its tipping point compared to another population, which is lying on a shallower potential (less stable) (Fig. 2). Our findings also suggested that the utility of EWS was strongly determined by factors that are intrinsic to a population such as genetic variation, net reproductive rate and plasticity.

Higher strength of plasticity in the phenotype significantly affected the detectability of population collapse (Fig. 2). Strength of plasticity altered the response and the dynamics of a population to changes in the external environment. When plasticity was high the response of the phenotype tracked the environment perfectly causing the population to stabilize in response to fluctuating changes in the environment (Fig. S3) (Charmantier et al., 2008; Reed, Schindler, & Waples, 2011). The stability of a population is also reflected in short return time to equilibrium (van Nes & Scheffer, 2007). This was true for the case where the strength of plasticity was high (Fig. S1). Moreover, the basin of attraction for a highly plastic population was deeper when compared to a less plastic population, which was shallower. To push a highly plastic population from the deepest point in its effective potential energy curve to the tipping point, the energy loss would be more, compared to a less plastic population which was lying in a much shallower basin. The loss in energy from driving the highly plastic population from the deepest point to its tipping point was reflected in stronger EWS when one compared it with another less plastic population. Consequently, for the same rate of change in the environment, higher values of plasticity lead to stronger and more reliable EWS of population collapse.

Similarly, the effective potential of our dynamical system was also altered by genetic variation (Fig. 2), with higher genetic variation leading to a shallower stability landscape than that of a population with lower genetic variation. High additive genetic variance created what was called a ‘genetic load’ that lowered the overall population growth rate (Lande 1976; Lande and Shannon 1996; Chevin, Lande, and Mace 2010). This was due to stabilizing selection acting on the mean phenotype. Due to this genetic load caused by high additive genetic variance, population growth rate at equilibrium was lower causing the effective potential to be closer towards the tipping point compared to another population having a lower genetic variation. Hence for the same rate of external environmental change, the loss of potential energy was different when one compared a population with high genetic variation to that with low genetic variation. As a result, the performance of EWS was affected with slightly lower detectability and lower reliability of population collapse when there was higher genetic variation (Fig. 3, Fig. 5B). This result however did not mean that genetic variation was detrimental for long-term population persistence. Time to extinction was however longer for populations with higher genetic variation when compared with a population which had a lower genetic variation in the trait (Fig S2)(Robinson, Wares, & Drake, 2013; Willi & Hoffmann, 2009). High genetic variation is necessary for rapid adaptation via genetic evolution, which then aids in evolutionary rescue (Ashander, Chevin and Baskett, 2016). However, our model simulations and analyses of EWS were mainly focused on short transient dynamics after the start of the external optimum change. Further, due to continuous and fast rate of environmental deterioration, adaptation or evolutionary rescue by genetic evolution (Lande 2009; Chevin and Lande 2010) was not possible in our model and the populations eventually declined.

Decreasing population size in response to environmental deterioration and directional selection acting towards a mean phenotype could both lead to depletion of additive genetic variance (Barton & Keightley, 2002). We took this account in our simulations by following a framework developed in (Burger, 2000) (see appendix 5). Using this framework, we find that depletion of genetic variation over time did not affect the strength of EWS compared with when genetic variation remained constant (See Fig S6).

Net reproductive rate also had a significant effect on the stability of populations, detectability and reliability of EWS of population collapse. Higher net reproductive rate buffered the population from declining immediately after the environment shifted when compared to a population with lower reproductive rate (Juan-Jordá et al., 2015). Consequently, for the same rate of change in the external optimum environment, populations with lower reproductive rate recovered slowly due to a wide basin of attraction than populations with higher net reproductive rate (Fig. 2, Fig. S1) and hence strength in EWS was more pronounced in the latter.

Our modeling results suggested that including fitness-related trait information could significantly improve the strength and reliability in EWS of population collapse. Trait-based EWS (*SD+trait*) produced strongest and reliable signals. Inclusion of phenotypic trait information alongside the leading indicators produced less distributed, consistent and stronger Kendall’s tau values compared to the generic abundance-based EWS (Fig. 4, Fig S7-S8). This is because in response to a directional change in the external optimum environment, shift in the mean phenotypic trait occurred over time even if there were no significant increases in either AR1 or SD. Furthermore, including information from the mean phenotype alongside EWS statistics would depend not only on whether the mean phenotype responded to the external forcing but also on the nature of the external environmental forcing itself (Clements & Ozgul, 2016b). Body size is suggested to be an ideal candidate to be included with the leading indicators (Clements & Ozgul, 2016b). Shifts in body size have previously been shown to occur before population collapse in whales, with true positive EWS being detectable earlier when information from the body size was included in the EWS (Clements et al., 2017). Furthermore, it has been suggested that in certain environmental change scenarios, it is possible for body size to shift before a population decline and thus act as a warning signal (Baruah *et al.*, 2018 *(in press)*).

In conclusion, we showed that predictability of population decline was affected by factors intrinsic to a population like genetic variation, strength in adaptive plasticity and net reproductive rate. Changes in these factors can not only alter the underlying phenotypic and population dynamics, but also the effective potential landscape. Consequently, due to the alteration of the effective potential landscape, the detectability of EWS was hampered. However, if information from fitness-related trait was incorporated with EWS, the strength in predicting population collapses increased significantly. Our results suggests that the inclusion of trait dynamic information alongside the generic EWS should provide more accurate forecasts of the future state of biological systems.

## 5. Code availability

R version 3.5.1. R script for the code is made available: https://github.com/GauravKBaruah/Eco-Evo-EWS

## 6. Data availability

will be made available at Dryad repository.

## 7. Author contributions

GB conceived the ideas and designed the methodology; GB analyzed the data with feedback from CFC and AO; All authors contributed to the drafts and final approval for submission.

